# Dissociation of early and late face-related processes in Autism Spectrum Disorder and Williams syndrome

**DOI:** 10.1101/2021.04.07.438774

**Authors:** Alice Gomez, Guillaume Lio, Manuela Costa, Angela Sirigu, Caroline Demily

## Abstract

Williams syndrome (WS) and Autism Spectrum Disorders (ASD) are psychiatric conditions associated with atypical but opposite face-to-face interactions patterns: WS patients overly stare at others, ASD individuals escape eye contact. Whether these behaviors result from dissociable visual processes within the occipito-temporal pathways is unknown.

Using high-density electroencephalography, multivariate signal processing algorithms and a protocol designed to identify and extract evoked activities sensitive to facial cues, we investigated how WS (N=14), ASD (N=14) and neurotypical subjects (N=14) decode the information content of a face stimulus.

We found two neural components in neurotypical participants, both strongest when the eye region was projected onto the subject’s fovea, simulating a direct eye contact situation, and weakest over more distant regions, reaching a minimum when the focused region was outside the stimulus face. The first component peaks at 170ms, an early signal known to be implicated in low-level face features. The second is identified later, 260ms post-stimulus onset and is implicated in decoding salient face social cues.

Remarkably, both components were found distinctly impaired and preserved in WS and ASD. In WS, we could weakly decode the 170ms signal based on our regressor relative to facial features, probably due to their relatively poor ability to process faces’ morphology, while the late 260ms component was highly significant. The reverse pattern was observed in ASD participants who showed neurotypical like early 170ms evoked activity but impaired late evoked 260ms signal. Our study reveals a dissociation between WS and ASD patients and point at different neural origins for their social impairments.

## Introduction

Although Autistic Spectrum Disorder (ASD) and Williams syndrome (WS) are both complex developmental conditions with core behavioral deficits in social cognition and face processing, the nature of these deficits differs greatly. Whereas patients with ASD struggle with social interaction and making eye contact, patients with WS seek to do both.

ASD is a neuro-developmental disorder characterized by deficits in social communication and social reciprocity, as well as repetitive and stereotyped behaviors. Despite extensive research into the biological factors underlying its pathology, behavioral observations remain the principal method of diagnosis (American Psychiatric Association & Association, 2013). Social deficits, most notably a failure to attend preferentially to the eyes of others, are signs of autism that are observable as early as the first months of life (Jones & Klin, 2013; Warren Jones et al., 2008).

WS is a rare neurodevelopmental disorder caused by a hemizygous deletion of approximately 25 genes on the 7q11.23 chromosomal region (Korenberg et al., 2000) resulting in a phenotype comprised of medical, cognitive, affective, and neurophysiological impairments (Bellugi et al., 2007). A core behavioral component of this syndrome is increased motivation for social interaction with an apparent lack of fear of strangers (Doyle et al., 2004; Zitzer-Comfort et al., 2007). Patients with WS have been described as acting as if “everybody in the world is their friend” (Doyle et al., 2004) and their appetitive drive for social interaction and social closeness with other people is commonly likened to hypersociability (Pavlova et al., 2016).

Patients with WS display atypical facial processing behaviors which are clearly distinct from those of patients with ASD. Specifically, during social interaction, patients with WS appear to exhibit “face fascination” (Jarvinen et al., 2015) from infancy (Mervis et al., 2003). Relative to more general forms of visuo-spatial processing, WS patients demonstrate comparative strengths in face processing (Bellugi et al., 2000; Paul et al., 2002). However, studies have also noted atypical processing in WS patients of eye and mouth regions of the face. Overall, face scanning patterns of individuals with WS differ from those produced by ASD patients. Patients with WS show an increased preference for eyes over the mouth region (Riby et al., 2009; Tager-Flusberg et al., 2003) in upright faces (Hirai et al., 2016). This pattern of dissociation in face processing between patients with ASD and patients with WS is intriguing, especially considering that face-to-face interaction is a critical ability on which future social interest and social skills are based (e.g., Ferrari, Paukner, Ionica, Suomi, 2009; Dettmer, Kaburu, Simpson, Paukner, Sclafani, Byers, Murphy, Miller, Marquez, Miller, Suomi, Ferrari, 2016; Farroni, Csibra, Simion, Johnson, 2002).

Faces are multidimensional visual stimuli offering a rich variety of information to observers (Lee et al., 2012). The multidimensional nature of this information is key to social interaction. Faces not only convey permanent and stable information such as gender (male or female), race (e.g., Chinese or Caucasian), identity (e.g., John or Mary), but also dynamic and transient information such as emotional expression, direction of attention, and intention (Lee et al., 1998).

The neurobiological substrate of these complex cognitive processes relies on at least three brain structures: the occipital and fusiform face areas and the posterior superior temporal sulcus (STS). Studies using PET-scan and fMRI techniques have showed convincing evidence that blood flow increased in the fusiform gyrus in responses to human faces compared to other stimulus viewing, thus leading to its labelling as the fusiform face area (FFA) (Haxby et al., 1994; Kanwisher et al., 1997; Kanwisher et al., 1998). Techniques with high temporal definition such as EEG have also revealed face-specific evoked potentials (ERPs) in the occipito-temporal cortex. Most notably the N170, a negative potential that peaks at 170 msec is elicited by faces (Allison, 1999) and by face components, especially the eyes (Bentin, 1996). Furthermore, evidence also suggests that downstream visual regions, such as the occipital face area, a relatively neglected region in face processing, may play a role in coding critical face-parts components (Gauthier et al., 2000; Pitcher, Walsh, et al., 2011) during the early stages of processing (Mattavelli et al., 2019; Sadeh et al., 2010). Studies have so far demonstrated that BOLD activity in occipital face area (OFA) correlates with ERP face-selectivity at 100–110ms after stimulus onset while BOLD activity in FFA correlates with ERP face-selectivity at 170ms (Sadeh et al., 2010). Hence, parts of the face are first processed in the OFA and then computed holistically in the FFA (Yovel, 2016).

Along the occipito-temporal visual stream the STS plays a key role in the human face-perception system, but it is also one of the key components of the ‘social brain’. Indeed, brain regions in and around the superior temporal sulcus of both hemispheres may be involved in the analysis of actual or implied facial movements and related cues that provide socially relevant information, such as emotional expression (Pitcher, Dilks, et al., 2011; Puce et al., 1998; Schobert et al., 2018; Winston et al., 2004) and gaze direction (Burra et al., 2017). Although establishing which cortical areas generate the N170 is problematic owing to source localization issues (Slotnick, 2004), electrophysiological studies suggest that the STS might be crucially involved in generating this ERP in response to face stimuli (Nguyen & Cunnington, 2014). Studies that have attempted to localize the N170 to face-selective regions suggest that the N170 reflects neural activity arising from the FFA (Horovitz et al. 2004), pSTS (Henson et al. 2003), or both the FFA and pSTS (Sadeh et al. 2010), but not the OFA.

The superior temporal sulcus has also been hypothesized as a key structure associated with the social-interaction deficits that are typical in patients with ASD (Allison et al., 2000; Dakin & Frith, 2005; Zilbovicius et al., 2006). Specifically, this hypothesis points to the STS as playing a role in the early stages of visual processing analysis of social cues. A link between perceptual abnormalities and specific disturbances in social cognition associated with autism has been proposed. Anatomical and functional alterations in the STS region of ASD individuals corroborate this hypothesis. PET-scan studies have described localized bilateral temporal hypoperfusion in children with autism (Zilbovicius et al., 2000) as well as a decrease of grey matter and an elongation of the superior temporal sulcus (Boddaert et al., 2004; Hotier et al., 2017). In a high-density EEG study while participants performed a face perception task, Lio and colleagues (2021) recently showed that evoked activity in the STS occurring at 240 ms after stimulus onset was significantly high in neurotypical subjects but weak in ASD patients. Furthermore, using this specific activity evoked in the STS, these authors were able to identify ASD patients and neurotypical subjects with machine learning algorithm with a relatively high degree of accuracy (>80%). This suggests that STS activity is crucial for coding the social value of faces. To measure the possible modulation of the STS signal for face subcomponents (eyes, mouth, eyebrows, cheeks etc), Lio *et al* (2021), in a second task, constrained subjects’ attention by presenting each face region in the foveal field. With this design they showed that, in the neurotypical population, the evoked activity is eyes sensitive, that is, the neural signal in the STS source is higher when subjects focused attention on the eyes. In contrast, such modulation was not found in patients with ASD, where evoked activity is not further enhanced when perceiving the eyes. These findings thus suggest that in the neurotypical population the 240ms evoked response in the STS is related to the social meaning of faces and more specifically gaze sensitive. Such late gaze-sensitive activity appears to be lacking in the ASD population.

Although, from the perspective of social cognition and face processing at least, ASD and WS conditions appear to reflect two sides of the same coin, the neural substrates that have been proposed as involved in WS diverge from those thought to be implicated in ASD. The regions mainly associated with motivational value (amygdala) and executive control (frontal lobe) have been postulated to play a role in the observed cognitive abnormalities characteristic of WS. It has been hypothesized that maladaptive social behavior in WS is due to lack of inhibition and impaired emotional processing consequent to frontal (Porter et al., 2007) and amygdala dysfunction (Bellugi et al., 1999; Meyer-Lindenberg & Hariri, 2005; Reiss et al., 2004).

Other studies, however, have suggested that the strong tendency demonstrated by WS patients to approach others and, in particular, to attend to faces (Jones et al., 2000) could be explained by additional dysfunctions in another set of neural substrates involved in visual processing. Indeed, WS patients show abnormalities in both the FFA and STS. The FFA has been found to be enlarged (Golarai et al., 2010) as well as structurally altered (Reiss et al., 2004) in these patients. Moreover, although WS patients show an overall reduction in brain volume (Chiang et al., 2007; Fung et al., 2012; Sampaio et al., 2008), cortical thickness of the superior temporal gyrus has been found increased compared to the neurotypical population (Green et al., 2016). Also, a recent fMRI study using a facial emotion matching task, has shown greater activation of the superior temporal sulcus in WS patients compared to patients with anxiety disorders and less activation in the occipital face area compared to controls (Binelli et al., 2016).

Electrophysiological studies also demonstrated that binding of face features in WS is atypical (Mobbs et al., 2004). In an ERP face recognition task, contrary to controls, who showed an N320 signal for upright face recognition and a P600 for inverted faces, WS patients produced a similar match-mismatch effect on N320 for both upright and inverted faces (Mills et al., 2000). In this same study, WS patients showed an abnormal increase in N170 amplitude when viewing trustworthy faces, suggesting that WS patients’ hypersociability may be linked to such enhanced early visual brain activity. In accordance with this idea, Shore and colleagues(2017a) have proposed that impairments in low-level perceptual processes might have cascading effects on social cognition.

In the present study, using the face perception task developed by Lio *et al*.(2021), we asked whether the atypical neural signal observed in patients with ASD in the STS at 240ms might also be observed in WS or whether there is a neural signature modulated by facial cues that reflects the behavioural dissociation between these two groups. Overall, we sought to determine whether the opposite nature of face-related behaviors observed in WS and ASD can also be reflected at the neural level.

We used the behavioral procedure (see Figure 1A) designed by Lio *et al*. (2021) that combines a face gender discrimination task with high density EEG recording. In this task, we were able to control the area of the face that subjects focus on and relate it to the cortical response. We examined the EEG brain activity of WS patients compared with data from a group of neurotypical subjects and a group of patients with ASD previously reported by Lio *et al*. (2021).

**Figure 1:**
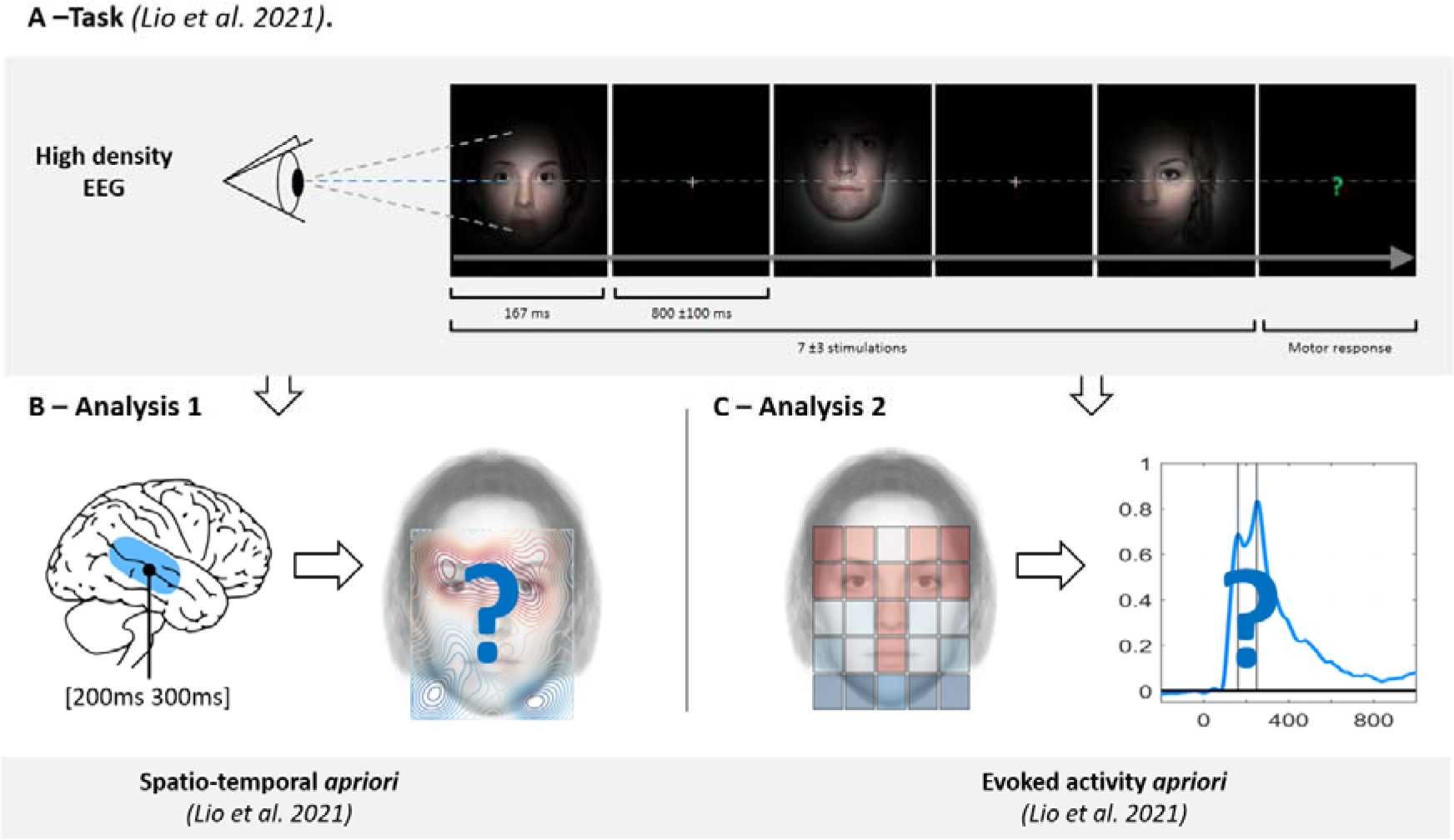
Methodological procedure summary. (**A**) **Facial-cue task** Participants were instructed to focus on a fixation cross and were then presented with a face stimulus masked by a Gaussian apodization window centered around the fixation cross (FWHM = 10°). The focused face area at the center of the screen was randomly drawn from a uniform distribution among 25 predetermined locations. A question mark was presented at random points (every 7±3 trials, at which point, participants were required to determine the gender of the last face stimulus that was viewed (left/right button press). (**B**) **Analysis 1.** The first analysis used a strong spatio-temporal a priori to build a map of cortical reactivity according to the face area that was focused on by each participant. Cortical reactivity was evoked in the region of the STS occurring 200-300ms after stimulus onset, as in Lio et al. The purpose of this analysis was to determine whether the neurotypical response to eye contact in this source was preserved in WS. (**C**) **Analysis 2.** The second analysis aimed to identify changes in cortical reactivity associated with a “facial cue” regressor over time (built from the Lio et al. Exp .2 study, see method), and to identify when in time EEG activity can decode the “facial cue” map. This spatial regressor predicts a maximum evoked activity when participants focus on facial cues, with a progressive decay in response to other parts of the face and a minimum activity outside the face, as in the neurotypical population (See Figure 3A for details).

Two sets of EEG analysis were performed. First, in a spatio-temporal a priori analysis we built a spatial filter from the STS source identified at 240ms by Lio *et al*. (2021) and used it to extract evoked single-trial activity. Then, for each participant, a map was generated indicating the level of evoked activity as a function of each viewed face area. We expected that, unlike what was found in ASD patients by Lio *et al*. (2021), face parts that carry a rich amount of social information, such as the eyes, will yield selective STS responses in WS patients, just as was found.

Second, we performed a single trial analysis with an evoked activity a priori. Using a “facial cue” map as regressor of evoked activity, we investigated how much the multichannel EEG signal was able to decode this predicted pattern of evoked activity at every time point. This analysis has the capacity to indicate at which point in time socially relevant facial features of the regressor induce cortical reactivity.

## Materials and methods

### Participants

We recruited 14 neurotypical participants (5 men and 9 women, mean age = 11.6, range = 6–21) and 14 WS patients (5 men and 9 women, mean age=11.6, range= 6-21) matched for age and gender. The number of participants in each group was selected to provide a balanced designed across groups, therefore we matched the size of groups to that acquired by Lio et al (2021). Although a larger sample size allows to find a smaller statistically significant difference, the difference found may not be clinically and scientifically meaningful, and furthermore, with respect to patients, we did not wish to submit unnecessary subjects to the procedure (Friston, 2012). Importantly, as shown by Lio et al., the EEG procedure was meant to be sensitive at the single subject level. We compared the present dataset with the EEG data of 14 ASD patients (14 men, mean age = 20, range = 18–21) recruited in a previous study by Lio et al. (2021, Exp. 3). All participants had normal or corrected to normal vision and all neurotypical participants had no history of psychiatric or neurological disease.

Participants were recruited through national advertisements from WS associations and from the Reference for Rare Diseases XXX Center (XXX hospital, XXX). Each patient received a diagnosis of WS following genetic assessment (deletion at 7q11.23) by the psychiatrist (XXX) involved in the study. Patients with WS and neurotypical subjects both participated in a short neuropsychological evaluation to assess visuospatial reasoning, logical thinking and verbal skills (matrix and similarities subtest, WISC-V Weschler, 2005) and visuospatial and auditory attention (Arrows and auditory attention subtest, NEPSY, Korkman et al., 2003). Unsurprisingly, patients with WS showed lower performances compared to controls in all of these tests (see Table 1). These subtests were selected because they strongly correlate with the general IQ (WISC-V, Weschler, 2005), and thus, allow us to determine the intellectual deficits of each patients with WS. As expected, scores from the control group were in the normal range. Neuropsychological data of ASD patients obtained from Lio et al.’s study involved IQ evaluation (WAIS IV or WAIS V). Patients with ASD showed normal intellectual abilities (Mean IQ= 97.20, SD= 27) with the exception of two patients showing lower scores on these tests (IQ < 75).

**Table 1:**
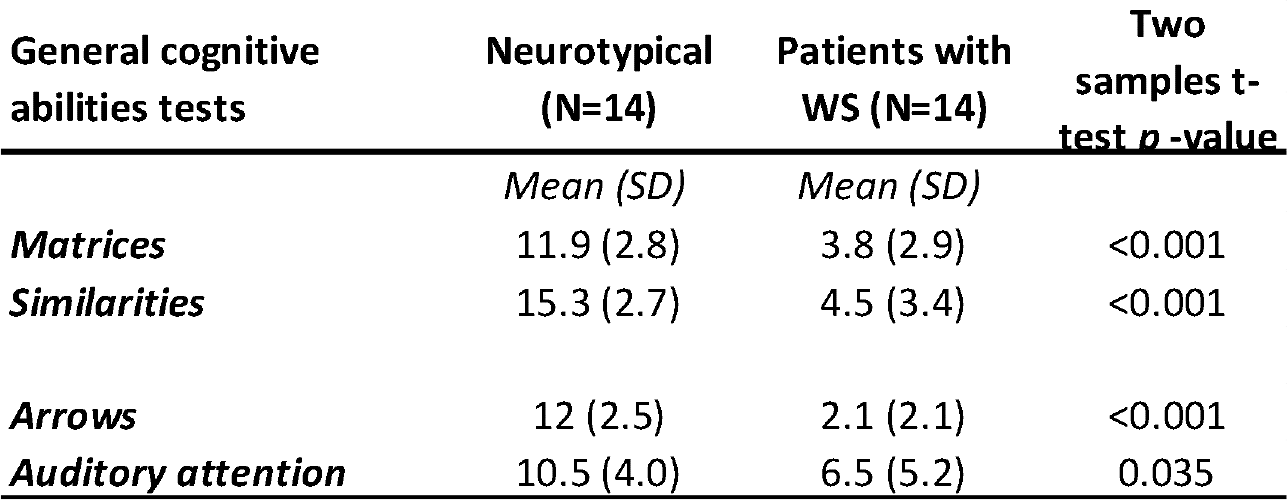
General cognitive abilities assessed in neurotypical participants and patients with WS.

The ability to recognize facial cues such as those related to emotional states is known to improve with chronological age. Significant changes in this ability have been found to occur in the first two years of life (for a review, Golarai et al., 2006) and to have a strong influence on social experience (Geangu et al., 2016). Chronological age, however, does not correlate with social approach ratings in Williams syndrome (Porter et al., 2007). Changes in the neural network dedicated to face processing are nevertheless related to chronological age (e.g., for the FFA, Nordt, Semmelmann, Genç, Weigelet, 2018; for the STS, Alaerts, Nayar, Kelly, Raithel, Milham, Di Martino, 2015).

Here, we used chronological age as a control basis for the amount of face-to-face experience participants accumulated and as an index of expected maturation processes in the face processing network. Experimental protocols based on intellectually age-matched designs are indeed known to be limited when dealing with population with intellectual disability, especially those with an heterogeneous profiles such as WS (Jarrold & Brock, 2004). Although WS patients are matched on global intellectual abilities, they are known to face more challenge in the visuospatial domain than in the verbal domain compared to other neurodevelopmental disorders. Therefore, we decided to control for the effect of chronological age by comparing patients with WS with an additional group of age-matched neurotypical participants and to statistically assess the existence of a mental age bias in our neurotypical population. To do so, we asked whether the intellectual abilities and chronological age of neurotypical participants (assessed with the matrix and similarities subtests) contribute to the observed effects in the EEG activity (see the section “Prediction of EEG decoding with behavioural measures”).

The study was approved by the French Sud-Ouest Lyon Bérard ethical committee (project N ° 2018-A02037-48). All methods were carried out in accordance with relevant guidelines and regulations. Prior inclusion, all participants and/or their legal representative provided written informed consent to participate in the study.

### Procedure

Participants were instructed to focus on a fixation cross (with a duration jittered 800 ± 100ms), which was followed by a face stimulus masked by a Gaussian apodization window (a Full Width at Half Maximum (FWHM)= 10°) centered on the fixation cross that appeared for 167ms (Figure 1A). Using this procedure, we were able to control which face region subjects were focusing on and the luminance distribution projected on to the retina. This procedure which reduces eye movements is similar to the “bubbles approach” method for studying the unit of face information processing (Gosselin & Schyns, 2001). To maintain subjects’ attention, every 7 (± 3) trial would take the form of a question mark which was presented on the screen instead of a face stimulus. Upon presentation of these trials, participants were required to recall the gender of the last face displayed and to respond using a button press with either their index or middle finger. The focus was on accuracy rather than speed as behavioural response was not the main goal of the study. Experimental testing was made up of three sessions each consisting of 500 trials. This was preceded by a training session with 100 additional trials to ensure that the gender discrimination task was performed above chance-level and that all participants understood the instructions. We set out to control the area of the face focused on by the participant in order to link it to cortical reactivity occurring at 200-300ms. Furthermore, we presented a limited number of trials to participants to ensure that the duration of task performance did not exceed an hour. This ensured that the task would be more amenable to the attentional abilities of young participants and patients with WS. Following pilot testing with adult participants with WS, we selected a total of 1500 face presentation trials (plus the 100 training trials).

### Stimuli

Face stimuli were identical to those used by Lio et al. (2021, Exp. 3) and delivered using Matlab and the Matlab Psychtoolbox. Each stimulus consisted of a neutral natural face controlled in proportion, position and luminance distribution. The faces are displayed on the screen at different locations on a grid, drawn from a uniform distribution. Then, each stimulus is multiplied by a Gaussian apodization window centered on the fixation cross. With this procedure, each trial consists of projecting different parts of the face onto the subject’s fovea while controlling the luminance distribution of each stimulus. The width of the aperture window was chosen so that it was large enough (Full Widths at Half Maximum =10° of visual angle) to make the recognition of gender or the identity of each stimulus easy across all trials (Figure 1A). Considering, the coordinate [0°; 0°] located between the two pupils of face pictures, each observed region is drawn from a uniform distribution on a [-8°; +8°] visual angle width, [-12°; +8°] height rectangle encompassing the whole face area. Thus, with this method, each region of the face is observed multiple times by the subject. In this study, in order to optimize the statistical power of the analysis given the reduced number of trials (1500), the region of focus for each stimulus presentation was no longer randomly selected from all of the pixels in the picture (20° height x 16° width, as in Lio. 2021, exp 2). Instead, regions were centered on one out of 25 rectangles defined as ROIs. These ROIs were generated by dividing the overall area of the studied face stimuli into a grid of 5x5, each ROI took the form of a rectangle (size, height = 4° x width = 3.2°). Given that we presented and recorded 1500 trials, the sampling density of face stimuli was 1500/25 = 60 trials / ROI / participant.

### EEG recording and preprocessing

We used the Brain Product™ actiCHamp system to record the electroencephalographic signal from 128 active electrodes (actiCAP 128Ch Standard-2) mounted in an elastic cap at 10-10 and 10-5 system standard locations (Oostenveld & Praamstra, 2001). All electrode impedances were kept below 50 kOhms. Subjects were seated in a darkened, shielded room with their head position controlled by an ophthalmic chin-rest device so that eye-level was aligned with the fixation cross. EEG data were recorded at a sampling rate of 5000 Hz with an online reference at the Fz electrode. Offline, data were band pass filtered using zero-phase Chebychev type II filters (Low pass - cutting frequency: 45 Hz, transition band width: 2 Hz, attenuation: 80 dB; order: 35, sections: 18 | High pass – cutting frequency: 0.3 Hz, transition band width: 0.2 Hz, attenuation: 80 dB; order: 9, sections: 5) and re-referenced to a common average. Next, data were epoched from 200 ms before to 400 ms after the stimulus onset.

Traditional EEG analysis considers the time course of individual channels. However, with modern high spatial density EEG (128 electrodes), we used multivariate signal processing algorithms as it can linearly combine channels to generate and aggregate representation of the data. This linear projection combines the information from the multiple sensors (128) into a single channel whose time course can be analyzed with conventional methods (temporal filtering, trial averaging), thus improving the spatial resolution and signal to noise ratio compared to traditional analysis (Parra et al., 2003, 2005).

### Analysis 1: Spatio-temporal a priori

First, a spatial filter allowed the extraction of single-trial activity evoked between 200-300ms in the STS (Figure 1B). This filter relies on the scalp topography of the source as established by Lio *et al*. (2021) (See section below for more details). Then, for each participant, a map was estimated of the level of evoked activity as a function of the focused face area. Finally, for each group, an average was generated that estimated which face area evoked the most activity.

### Single trial spatial filtering

Using a Group Blind Source Separation (gBSS) analysis and cluster-permutation test Lio *et al*. (2021), revealed a significant source in neurotypical participants consisting in a large evoked activity with a maximum at 240ms after the stimulus onset, and a source localization showing a maximum of activity around the lateral fissure (MNI coordinates = X: +65 -65 Y: -20 Z: 10) and a local maximum around the inferior temporal sulcus (MNI coordinates = X:+60 -60 Y: -40 Z: -20).

Based on the scalp topography of this source, which occurs between 200 and 300ms, we use spatial filtering to extract single-trial evoked activity in all participants. More specifically, to measure the evoked activity of the component identified in the Lio et al. (2021) study, we calculated a spatial filter for each trial using minimum variance beamforming (Van Veen et al., 1997; Vrba & Robinson, 2001) in combination with the spatial information estimated at the group level in the original study using gBSS (group Blind Source Separation) (Lio et al.. 2021, see also Albares et al., 2014 for a detailed description of the method).

For each participant, a map was generated indicating the relative level of evoked activity as a function of each of the 25 viewed face areas. To do so, we first average the evoked activity between 200ms and 300ms at each location, then we applied a Z-transform of the 25 obtained values in order to highlight the ‘most positive’ and the ‘most negative’ areas (See Figure 2A). This analysis allowed us to visualize for each participant how cortical sensitivity in the STS occurring at 200-300ms is affected by different face regions.

**Figure 2:**
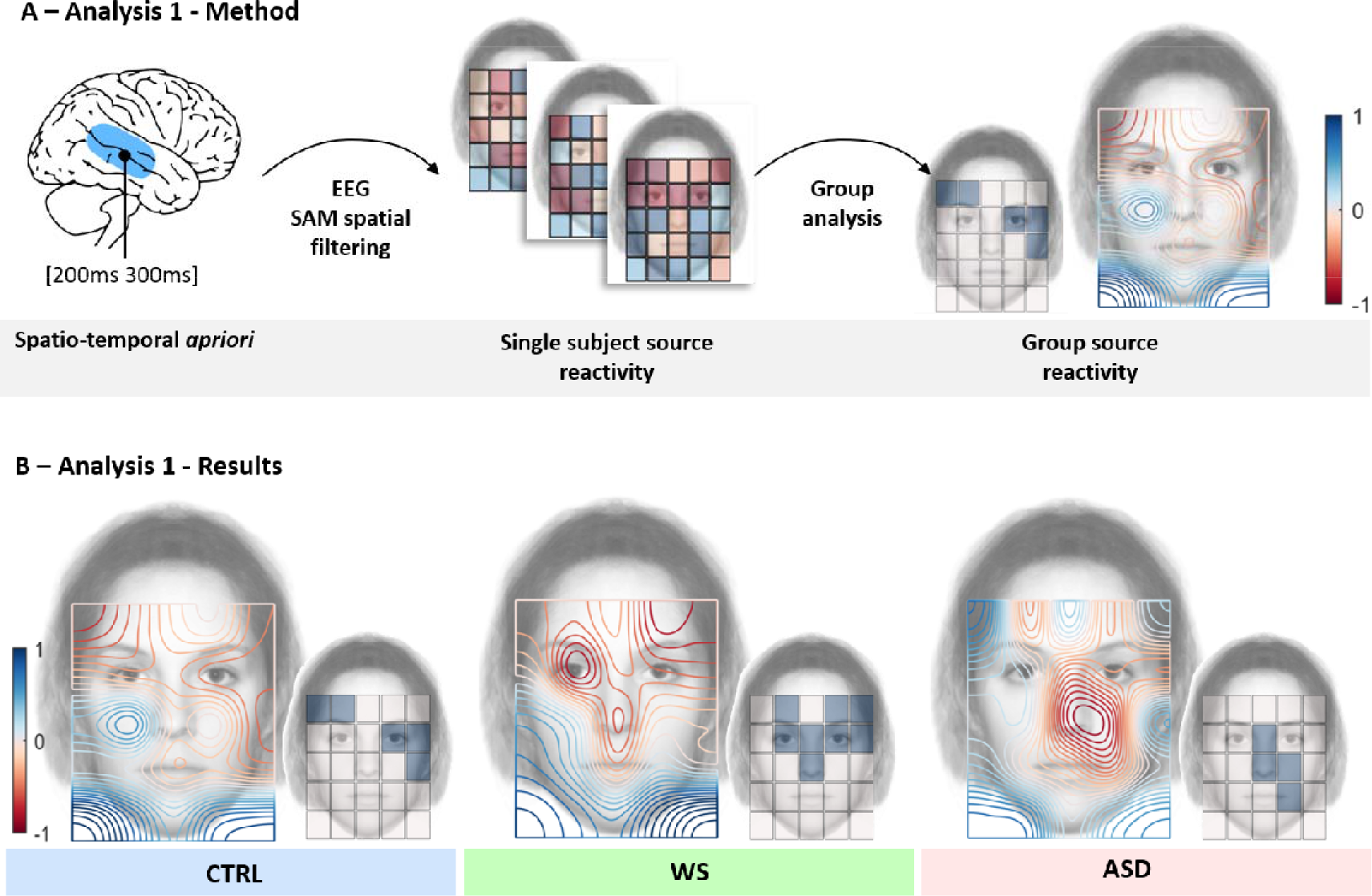
Method and results of the single trial evoked activity analysis. (A) Signal processing pipeline of this analysis 1. First, a spatial-filter was built to extract single-trial activity evoked between 200ms and 300ms in the superior temporal region at the single subject and group-level. **(B) Results of evoked activity mapped over a face area for the neurotypical participants (CTRL, in blue, on the left), patients with Williams-Beuren syndrome (WS, in green, in the middle); and patients with autistic spectrum disorders (ASD, in red, on the right).** For each group, the large face on the left (with a Red-Blue color scheme) shows a contour plot of interpolated results of the degree of evoked activity in the STS measured as a function of the face region attended to by participants. The small face on the right (with blue tiles) shows tiles which represent maximum and significant (p<0.05 FWER corrected) areas of evoked activity. Activity in young neurotypicals confirms that activity was eye-sensitive (significant on the left eye), as found previously in adults (Lio et al. Exp. 2). This was maximal in the upper part of the face, over the eyes and eyebrows, and decreased to reach a local minimum in the left and right lower corners, outside the face area. WS patients showed a similar activation map with significant activity over the eyes, eyebrows and nose region. ASD showed an atypical pattern with significant activity on the nose region and cheek.

### Group-level

Finally, we studied in each group of 14 subjects, which face region evoked a significantly the more pronounced activity by performing group statistical analyses on the 25 locations (25 non-parametric, N=14, one tailed, sign tests, p<0.05, FWER controlled using the maxT/minP multiple testing procedure (Westfall and Young, 1993, Figure 2). This process led to a statistical non-parametric mapping of the evoked activity in the superior temporal region (Figure 2B) for each group. For visualization purposes only, we processed a smoothed representation of the results obtained with an original resolution of 5x5 using bicubic interpolation of single-subject results and averaging interpolated maps at the group level (Figure 2B).

### Analysis 2: Evoked activity a priori

First, we generated a “facial cue” map regressor from neurotypical group data provided by Lio *et al*. (2021). Then, for each participant, we applied a multiple linear regression model to assess how much the multichannel EEG activity was able to decode the “facial cue map” over time (Figure 1C).

### Facial cue map regressor

We generated the “facial cue” map regressor by applying a sagittal symmetry and subsampling the cortical sensitivity map generated by neurotypical participants as in Lio et al.’s study (2021) (See Figure 3A left). The obtained regressor implies that evoked activity is maximal in the eyes and eyebrows region and gradually decrease over the nose and mouth and other face regions to reach a minimum outside the face.

**Figure 3:**
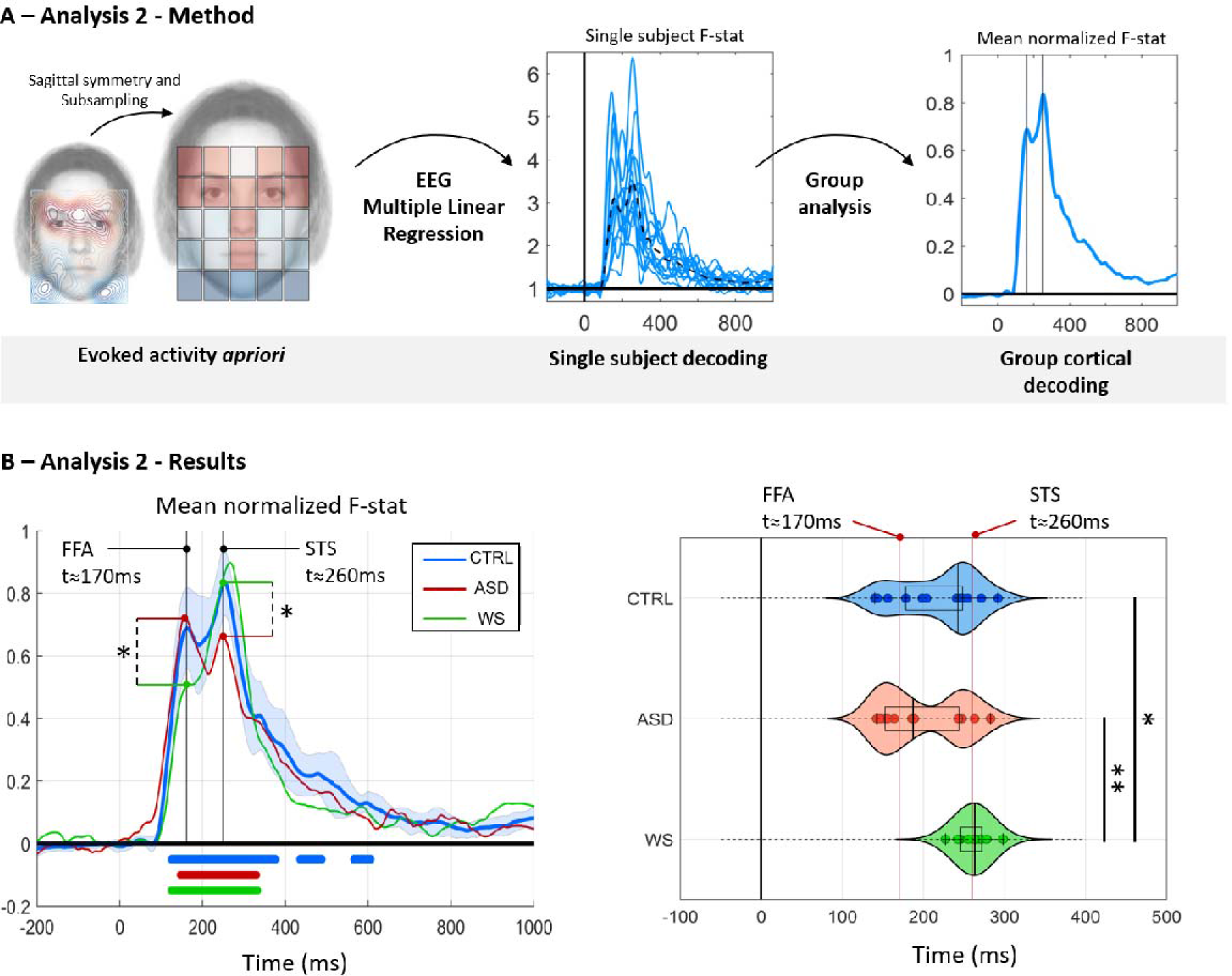
Prediction of evoked activity with face cue map spatial regressor. (A) Signal processing pipeline for the second analysis. The ‘socially relevant face cue map’ regressor was generated from the neurotypical evoked activity (applying a sagittal symmetry and subsampling) (left part). Then, a multiple linear regression is applied and the Fischer statistic denoting the quality of decoding obtained at each time point between 0-1000ms is generated for each subject (see dotted lines, on the middle graph). Finally, each decoding signal (F-stat) scaled between zero (min) and one (max) for each subject was averaged for the group analysis (right graph). **(B) Results from decoding the facial-cue map for the neurotypical participants (CTRL, in blue), patients with Williams-Beuren syndrome (WS, in green) and patients with autistic spectrum disorders (ASD, in red).** On the left graph, F score for the decoding of each group as a function of time. Significant F values are underlined in blue, green and red for neurotypical, patients with WS and Patients with ASD respectively. At the group level, the neurotypical decoding curve (blue) presents two marked peaks, one at 170ms post stimulus onset, the second at 260ms post stimulus onset. The ASD population (red curve) produced a significantly higher decoding peak than the WS population (green curve) at the earlier timing (170ms – p<0.05 FWER corrected). A reversed pattern was found at the latter decoding peak (260ms post-stimulus onset, WS>ASD p<0.05 FWER corrected). On the right graph, the distribution of the peak for each individual in each group is plotted over time. The distribution in the neurotypical population (blue) and in patients with ASD (red) is bimodal with a dominance at 260ms for neurotypical participants and at 170ms for patients with ASD. The distribution in WS patients is strictly unimodal.

### Multiple linear regression of EEG activity

With this analysis, we avoided making any spatial modelling assumptions with regard to sources or anatomy and relied entirely on the statistics of the observed data and its covariation with observable stimuli (Parra, Spence, Gerson, Sajda, 2005). Here, we denote **x**(*t*) as the vector of multidimensional EEG data (from all the 128 channels from the recording) at time *t*. A linear projection combines the information from the multiple sensors (128) into a single channel **y**(t), whose time course can be analysed with conventional methods (Parra, Spence, Gerson, Sajda, 2005). A vector w(t) is selected or calculated based on constraints or desired attributes of the time series **y**(t). Her, we aimed to find a weighted matrix w(t), at each time points, that could discriminate at the single-trial the level of evoked activity expected for the part of the face presented.

A multiple linear regression model was calculated at every time point, to predict the facial cue map regressor Y based on the multichannel EEG activity **x** (t): **Y**= w(t) * **x** (t) + n(t). Significant regression equation is reported using the F statistic, at every time point (between 0-1000ms), which represents how the quality of the decoding signal or how the EEG evoked activity at t codes for the facial cue map regressor. The F statistic is scaled between zero (min) and one (max) for each subject and averaged for the group analysis.

For each individual, we collected the timing peak of the F statistic. We tested if the temporal distribution of the peak differed across groups (Neurotypical, ASD, WS) using a Kruskall- Wallis test. To account for potential behavioural confounding, we performed a multiple linear regression to determine whether the variability of the onset of peak decoding was dependent on behavioural variables. Specifically, we performed a multiple linear regression (stepwise) to predict when peak decoding occurs as a function of task accuracy, visuospatial reasoning ability (matrix), linguistic reasoning ability (similarities) and age. In ASD patients, the linear regression was performed using the verbal and the Performance IQ as regressor, instead of the visuospatial and language reasoning abilities scores.

## Results

### Behavioral results

In the gender discrimination task, WS patients correctly identified 67% of faces while neurotypical subjects and ASD were able to reach 91% and 89% of correct responses, respectively. Although WS patients were less accurate at discriminating gender compared to the other groups (WS vs ASD: *t*(27) =5.25, *p*<0.001 and WS vs neurotypical: t(27)=-5.61, *p*<0.001) they were able to perform above chance-level (*t*(13)=4.17, *p*<0.001). Reduced gender discrimination performance in patients with WS may be explained by deficiencies in both general working memory and face processing capacities (Herwegen, 2015; Rhodes et al., 2010). The accuracy did not vary as a function of the face part (*F* (24, 975) = 0.7, *p*=0.85) and face parts did not interact with the group effect (*F* (48, 975) = 0.74, *p*=0.91).

### EEG results

#### Analysis 1: Spatio-temporal a priori

Group results are shown in Figure 2. We report the degree of evoked activity in the STS measured as a function of the face region attended by participants with maximum and significant (p<0.05 FWER corrected) areas of evoked activity. As reported by Lio et al. (2021) in neurotypical subjects, the evoked activity is eye-sensitive. These subjects produced a maximal evoked activity in the upper part of the face (p<0.05 FWER corrected, indicated by blue tiles over the right small face), over the eyes and eyebrows, and a local minimum in the left and right lower corners, outside the face area. These results replicate the findings of Lio et al. (2021) in a young neurotypical population and support the idea that face related signal in the STS is not strongly influenced by the age of participants (see Figure 2).

Although, the maximal evoked activity is above the left eye brow and at the right eye (and at the right from the right eye) which may suggest that a developmental trend could exist.

Patients with ASD showed an atypical pattern with significant activity on the nose region and cheek (p<0.05, FWER corrected, see Figure 2, bottom left). These previously reported results are consistent with neurotypical eye tracking behavior in face perception tasks whereby more attention is paid to the eye region than to the mouth areas as well as with previous eye-tracking studies of how ASD patients attend to faces (Hernandez et al., 2009; Riby et al., 2009).

WS patients showed an activation map with significant activity over the eyes, eyebrows and nose region (p<0.05, FWER corrected, see Figure 2, bottom center) and local minimum in the left and right lower corners and outside the face area. The pattern of evoked activity at 260ms in the STS contrast that of patients with ASD and matches that of neurotypical participants. This result is striking, because we show the existence of a neural process that appears to be preserved and robust in these patients even though the WS population we tested is both younger and has a lower IQ than patients with ASD in our sample.

#### Analysis 2: Evoked activity a priori

The maximal evoked activity for neurotypical participants (from Lio et al., 2021), from our younger population and from our patients with WS is reminiscent of a T-shape over the face: the eyebrows, eyes, nose and mouth (and these face parts carry rich source of social information), whereas we observe lower activity for other areas of the face that are considered to have little social salience. Consistently, the minimum of the activity was recorded when patients’ eyes were forced to look outside the face. We further built a face cue map spatial regressor generated by applying a sagittal symmetry and subsampling the cortical sensitivity map obtained by the neurotypical population and label this regressor a ‘face cue map’.

Then, for each subject a temporal signal of the Fischer statistic denoting the quality of decoding obtained between 0-1000ms was generated and compared across groups. We report each decoding signal (F-stat) scaled between zero (min) and one (max) for each subject and averaged for the group analysis in Figure 3.

At the group level, the F-stat value indicates that the decoding of the signal by the ‘face cue map’ was significant around the 150-350ms time range after stimulus onset in all three groups (*p*<0.05, FWER corrected, see Figure 3 bottom left). The decoding curve in neurotypical yielded two marked peaks, one appearing at 170ms after stimulus onset, and a second appearing at 260ms after stimulus onset. This suggests the presence of two independent and/or interacting processes that are dependent on facial cues.

The spatio-temporal dynamics of face processing have been well-studied using intracerebral electrophysiological recordings (e.g., Barbeau, et al. 2008). While late components (after 200ms) are diffuse in the brain (with peaks in various temporal areas) early components around 160-170ms have only been identified in the fusiform gyrus, around the FFA. We will therefore assume that the evoked activity modulated by facial features at 170ms arise from the FFA while the late component that was described by Lio et al. (2021) originates from the STS.

Although two response peaks with similar timing were identified in all groups, we observed different response level for these two processes for the two patients’ groups (See Figure 3A). The ASD population showed a significantly greater decoding peak than the WS population at the earlier timing (170ms post stimulus onset, ASD > WS – *p*<0.05 FWER corrected). A reversed pattern can be found at the latter decoding peak: patients with WS showed a significantly greater decoding peak than the ASD population at the later timing (260ms post-stimulus onset, WS>ASD *p*<0.05 FWER corrected). Because this method of EEG analysis requires no assumptions regarding cortical source, its use offers a significant advantage over more traditional alternatives, especially when applied to neurodevelopmental disorders.

For each group, we further tested which of the two peaks were maximal at the subject level. The temporal distribution of the maximum peak of decoding can be found in Figure 3B. These temporal distributions were found to differ between groups (Kruskall-Wallis, *Chi^2^*(2, 39) =11.9, *p*=0.0026). For neurotypical participants, we found that the distribution of the maximum peak of decoding had a bimodal distribution with a dominance for the second process. This distribution indicates that face decoding goes preferentially through a social decoding in neurotypical participants. ASD patients also showed a bimodal distribution but the early process at 170ms was significantly prominent. Consistent with behavioral results, this pattern suggests that faces’ related processes within the ventral regions are well-preserved. Finally, WS patients showed a strictly unimodal distribution such that activity was focused around the second process only.

Post-hoc analyses revealed a clear dissociation between both the WS and ASD patients (*p*<0.01 FWER corrected) and the WS group and neurotypical population (*p*<0.05 FWER corrected). This result combined with the low performance of WS patients in the gender discrimination task, suggests a preferential use of the dorsal face processing regions during face processing tasks compared to the ventral and is consistent with social cognition biases found in this syndrome.

### Prediction of EEG decoding with behavioural measures

A significant regression equation was found for neurotypical participants *F* (1,13) =8.56, *p*=0.013, with an R² of 0.42 to predict decoding timing. Visuospatial reasoning abilities (matrix score) were the only significant predictor of the maximal onset of decoding (*t*=-2.9, *p*=0.013). Chronological age and linguistic reasoning abilities (Similarities test) were not significant predictors. In patients, no significant regression equation was found, most likely due to the lack of variability across participants on the decoding timing in each sample.

## Discussion

The present study confirms the previous results reported by Lio *et al*.(2021) and extend these findings by dissociating the time course of neural processes involved in face perception in WS and ASD patients.

Our analyses confirm that activity in neurotypical participants can be evoked by viewing the socially salient region of a natural face stimulus such as the eyes. Furthermore, the evoked activity originates bilaterally in the superior temporal regions and peaks at 260ms after stimulus onset. Notably, we show that the evoked STS eye sensitive response is also present in the WS group, contrary to what is observed in ASD participants by Lio *et al*. (2021). This reveals a dissociation among these two patients’ groups and suggest that although both syndromes are associated with social disturbances, their impairment at the neural level may have a different origin.

When examining face processing related neural signals along the occipito-temporal stream, we found two peaks in neurotypical participants: the first at 170ms, an early signal known to be implicated in low-level face features, the second at 260ms, a late component implicated in decoding salient face social cues. Remarkably, both components were found distinctly impaired and preserved in WS and ASD. In WS, we could weakly decode the 170ms signal probably due to their relatively poor ability to process faces’ morphology while the late 260ms component shown to be eye sensitive was highly significant. The reverse pattern was observed in ASD participants who showed neurotypical like early 170ms evoked activity but impaired late evoked 260ms signal.

Hence, we report a functional double dissociation between early (170 ms) and late (260ms) neurophysiological responses to faces in two neurodevelopmental disorders known to offer dissociable and opposing behaviors in facial processing and social cognition. Face processing is thought to involve the ventral pathway (through the OFA and FFA) for low-level features that are generally invariant, such as gender, age and identity, while the dorsal pathway (through the STS) is thought to process facial movements related to emotional expressions, gaze cues or intentions (Bernstein & Yovel, 2015; Haxby et al., 2000; Pascalis et al., 2011). The latter is also considered to be the gateway to an extended system of social perception encompassing the orbitofrontal cortex and the amygdala (Truett Allison et al., 2000). Indeed, the present finding supports the hypothesis that early (170ms) visual processing of faces in the temporo-occipital area (FFA) and late social processing of faces in the temporal sulcus (i.e. STS) are functionally dissociable (Figure 4) as previously suggested using rTMS (Dzhelyova, Ellison, & Atkinson, 2011)

**Figure 4:**
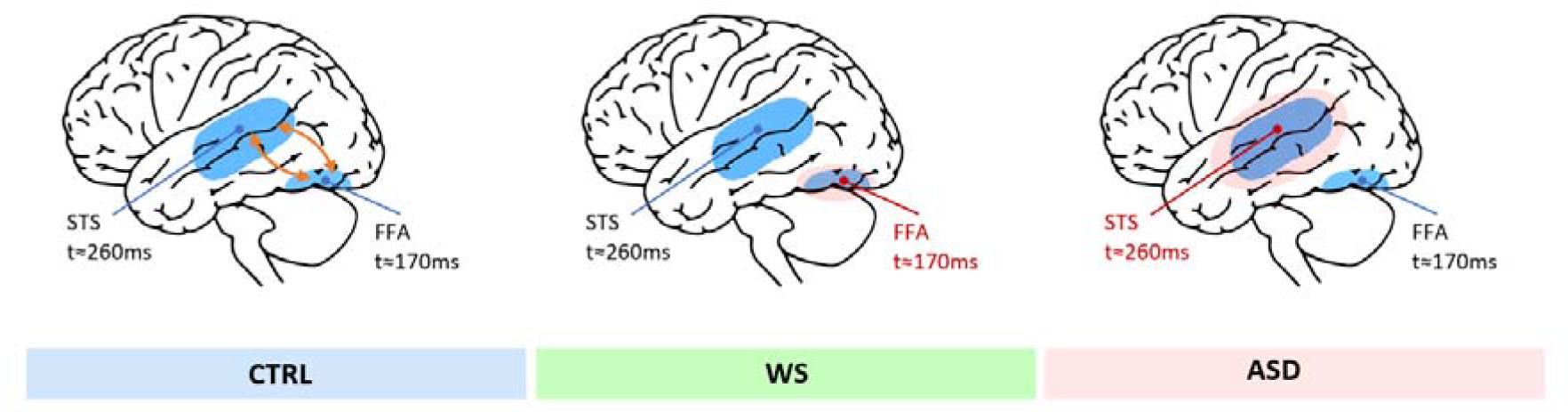
Time course of face processing along the temporo-occipital pathways in neurotypical subjects, WS and ASD. On the left, a schematic proposal of the two components interactions in face processing of neurotypical participants (blue): the early component (t=170ms) in the Fusiform face Area (FFA) and a late component (t=260ms) in the Superior Temporal Sulcus (STS). In the middle, a schematic representation displaying the proposed altered functioning (in red) of the early component of face processing (t=170ms) in patients with Williams syndrome (WS). On the right, a schematic representation displaying the proposed altered functioning (in red) of the late component (t=260ms) in patients with Autism Spectrum Disorder (ASD).

It could be argued that our study design may bias interpretations because patients with WS and patients with ASD present different intellectual abilities. However, we first report a preserved STS signal in the WS population at 240ms, and if mental age or chronological age had biased the results, one would have expected an opposite pattern (impaired WS patients, preserved ASD patients). Therefore, our results strikingly indicate that some high-level social processing of faces is well-preserved in patients with WS despite their low intellectual abilities and young age. Regarding the observed dissociation at 170ms, one could argue that the deficit showed in WS is due to their younger age compared to ASD patients. However, this interpretation is unlikely given that chronological age does not appear to have a significant impact on the onset of the decoding peak (in analysis 2) and that we replicated this result with the chronologically age-matched neurotypical participants.

The present data have theoretical implications for understanding WS and their characteristic atypical processing of social cues, particularly faces. Indeed, our findings suggest that patients with WS, when viewing the eyes of a face, lack specificity in their early neurophysiological response. Early neurophysiological responses to faces at 170ms are evoked by facial features (Allison, 1999; Bentin, 1996). This is thought to allow a consistent description of facial parts to be computed and processed in an holistic manner in the fusiform face area at 170ms (Gauthier et al., 2000; Pitcher et al., 2011; Yovel, 2016).

Previous fMRI studies have shown that, during face perception, less activity can be observed in patients with WS than in neurotypical participants in areas specialized in early processing of face parts (Binelli et al., 2016). Our data are consistent with these results and suggest that the FFA-driven process of combining parts of the face into a holistic representation may be specifically impaired in WS.

We show that early responses in patients with WS are not modulated by the type of facial features on display (i.e., eyes, eyebrow, mouth compared to other face-parts). Such an absence of modulation may suggest that patients with WS make little distinction between facial features and their possible relevance as a means for facial identification and subsequent holistic processing. Indeed, WS patients show increased activity at N170 amplitudes when viewing trustworthy versus untrustworthy faces (Mills et al., 2002; Shore et al., 2017).

In the present study, we applied blind source separation techniques to analyze high-resolution EEG data. EEG data were spatially processed to ensure that the source occurring at 170ms did not originate in the STS. We also show that WS patients produce robust responses to late processing of facial cues at 260ms in the STS. This observation is consistent with previous fMRI studies reporting greater STS activity in WS patients compared to patients with anxiety disorders while performing a facial perception task (Binelli et al., 2016).

Unlike the study by Shore et al. (2017), the altered neural response we found cannot be attributed to different patterns of eye fixation when looking at the face stimulus (Hirai et al., 2016, 2017; Mobbs et al., 2004). Indeed, our method assessed the neurophysiological response to specific regions of the face presented to the foveae of each participant. This experimental modulation is extremely relevant to WS, as it has been hypothesized that the facial scanning behaviour of these patients is influenced by a general difficulty in disengaging attention from salient targets, known as “sticky fixation“ (D’Souza, D’Souza, et al., 2015; Van Herwegen, 2015).

Our findings support two alternative interpretations. First, the role of the STS, which is typically implicated in late social processing of faces in neurotypical participants (Truett Allison et al., 2000), may be upregulated in patients with WS, thus leading to their characteristic highly social behavior. As a consequence, early analytical computations performed by the FFA (Kanwisher et al., 1998) would be bypassed, thus contributing to reduced performance in holistic face processing. Such interpretation would be coherent with reports of increased cortical thickness in the superior temporal gyrus of patients with WS (Green et al., 2016). Such interpretation would also fit well with the frontal dysfunction hypothesis, expressed as an inability to inhibit impulsive responses to social information (Porter et al., 2007) and the amygdala dysfunction hypothesis, expressed by atypical processing of emotional information relevant to social interactions (Bellugi et al., 1999; Meyer-Lindenberg & Hariri, 2005; Reiss et al., 2004). Children with WS are known to respond differently to social cues conveyed by the face (Doyle et al., 2004; Karmiloff-Smith et al., 1995, 2012; Riby & Hancock, 2009), and to process emotional cues atypically resulting in poor processing of social information from faces (Gomez et al., 2020). If the early analytical computation is by-passed it could explain why soft signs are not typically captured by patients with WS. Yet, in the present study, we do not evidence any atypical performance over the late component in patients with WS.

A second interpretation may be that early visual activity in temporo-occipital areas is dysfunctional in WS and are thus unable to discriminate specific facial features. This interpretation is supported by previous reports showing that the FFA is both enlarged (Golarai et al., 2010) and structurally altered (Reiss et al., 2004) in these patients. In our study, the poor decoding of the facial cue regressor is more likely related to their poor visuospatial abilities. In fact, in neurotypical participants the onset of the maximal decoding of the facial cue map regressor was predicted by visuospatial reasoning abilities (in the matrix subtest). That is, participants who showed poor visuospatial abilities where more likely to exhibit a late peak than those with good performance. This could also explain why WS patients showed a poorer performance on our gender discrimination task given their deficits in the perceptual holistic processes of faces. Therefore, it is possible that the first facial process in the ventral part of patients’ brain poorly decode facial features due to poor visuospatial abilities.

Subsequent cascading developmental consequences or compensatory strategies may induce a secondary upregulation of facial processing in other functional structures, namely the STS in this case, thus, leading to a concomitant increase in the social processing of faces. The role of cascading developmental consequences has already been theorized to explain atypical behaviors in other domains associated with Williams’s syndrome (such as mathematical impairments explained by their poor visuospatial abilities, Eckert et al., 2006). Such hypothesis is consistent with the atypical visual processing of faces from early infancy in WS (D’Souza, Cole, et al., 2015; Shore et al., 2017).

Overall, the notion of cascading pathways between early (170ms) and late (260ms) processes remains putative and could be coincidental. However, we argue that their development is closely intertwined. Still, whether the early neurophysiological face processing deficit in WS emerges from the neurodevelopmental consequence of the disorder or as the result of the indiscriminate social interaction exhibited by WS patients remains an unresolved question.

Given that patients with WS in our sample were relatively young (between 8 and 21 yo), it is possible that the atypical response at 170 ms might need more brain maturation and it may change with experience to become similar to that of neurotypical participants in older WS. However, such early adversity could induce long-term effects on visual processing or instead reach neurotypical performance later on, as is the case for face processing behaviour. Indeed, the frequency of pathological gazing at faces in WS individuals decreases after infancy and a more neurotypical focus on the eyes is observed with age (Martens et al., 2009).

One could see the lower intellectual abilities of patients with WS compared to patients with ASD and age-matched controls as a limitation, however, this ability are most representative from their disorder (i.e., intelligence range from 20 to 106, Ewart et al. 1993, with a remarkable deficit in visuospatial construction, Klein & Mervis 1999). Hence, we fully acknowledge that this factor contributes to the failure to decode the EEG signal based on the facial feature regressor at 170ms. Our aim was to identify what neurofunctional mechanisms could explain impaired face processing in patients with WS. Concerning IQ, most individuals with WS exhibit some degree of intellectual impairment, with the majority of adults scoring in the mild range of mental retardation on standardized intelligence tests, especially in the visuospatial domain (Howlin, Davies, Udwin, 1998). Therefore, from a clinical point of view, describing the impact of this neurodevelopmental syndrome, in the absence of mental retardation or visuospatial impairments, does not seem representative of the disorder itself. Nevertheless, to provide unbiased conclusions on the mechanisms, future studies should confirm that the rare subcategory of patients with WS having good visuospatial abilities also exhibit impaired face processing early decoding at 170ms. However, from a clinical point of view, the described findings are the most relevant as they apply to the majority of patients with WS.

### What are the interventional implications of these results? Targeting STS and FFA in neurofeedback for WS and ASD

Overall, our results raise the possibility of developing functional and behavioural rehabilitative procedures for both WS and ASD based on neurofeedback, a procedure in which self-regulation is stimulated by providing online feedback of neural activity to participants (Batail et al., 2019; Sitaram et al., 2017). By controlling specific neural substrates, it can be possible to pinpoint and modify specific behaviours. Due to its high spatial resolution, fMRI is usually used to target specific cerebral structures with neurofeedback (Linhartová et al., 2019; Okano et al., 2020). Most procedures using EEG trigger training in EEG signal coherence or frequency (Omejc et al., 2019). Rather than being based on modulation of disorder-specific biomarkers, most current EEG neurofeedback protocols are based on the modulation of a few spontaneous brain rhythms, mainly defined by the frequency of their oscillation (Batail et al., 2019; Sitaram et al., 2017). This strategy is widespread because spontaneous brain rhythms demonstrate a high signal-to-noise ratio in EEG recordings and can also be disrupted in some mental disorders (e.g., ADHD Lambez et al., 2020). However, within the broad field of psychiatric disorders, such methods lack specificity.

Here, ERPs could be used to feeding back levels of local cortical arousal back to patients with the purpose of improving the self-regulation of cortical excitability in the specific structure related to the disorder (Birbaumer, 1999). In order to respond to the specific needs of patients, training must be provided beforehand. Indeed, it has been demonstrated that patients with severe intellectual impairments, like some patients with ASD and WS, are able to follow training procedures to enhance the effectiveness of neurofeedback (LaMarca et al., 2018).

Our results provide unique and specific neurofunctional dissociation associated with face processing deficits in ASD and WS. In light of these findings, it may be beneficial to explore whether neurofeedback can help children with ASD or WS to enhance STS or FFA activity during face processing. Here, the goal would simply be to manipulate neural activity in the STS or FFA in order to rebalance social processing of face related to this structure, i.e., the identification of facial features. Evoked activity measured by high-density EEG while participants focus on different facial areas (either social, eyes, nose, brows, mouths or not) could be measured and used in real-time to display the decoding level from either the early (FFA, for WS) or late (STS, for ASD) component in real time, through visual, audio or other means, back to participants. The goal of this information would be to help participants self-regulate their cortical excitability related to face processing. Along with cognitive trainings, this procedure might assist patients in reassigning their attention to facial features for the purposes of identity recognition or social cognition (Lee et al., 2012).

One would expect such training to facilitate the identification of a neural strategy that can be employed by patients when viewing faces. Indeed, the idea would be to train patients both at the behavioural level to pay attention to socially relevant facial cues but to also to train them to engage their FFA and STS when viewing specific facial features (e.g., eye, eyebrows, mouth). The long-term goal could be that both groups of patients would engage the ventral and-dorsal areas in a more balanced way during social interactions.

To conclude, facial and social skills are essential abilities for navigating in our modern society. With the ongoing pandemic, the ability to decode social information from faces on a screen or from the eye region (as faces are partially occluded with masks) has recently become a critical tool in social interactions (Freud et al., 2020). This ability relies on a complex cerebral network (Yovel, 2016). Double dissociations using fine spatiotemporal analysis of this network provide essential evidence for understanding what, where and when neurocomputations are performed in our brain. In patients with Williams syndrome, poor decoding of facial features by low-level visual processing can result in some of the observed social symptoms that can have catastrophic influences on their lives. Future works may examine whether, in turn subtle changes in the balance of this network, through self-regulated neurofeedback training, can attenuate their characteristic social behaviour and related symptoms of social dysfunctions, such as anxiety.

